# A functional screen for optimization of short hairpin RNA biogenesis and RISC loading

**DOI:** 10.1101/2020.05.22.110924

**Authors:** Robert L Sons, Kyle W Kaufmann, Scott M Hammond

**Affiliations:** Department of Cell Biology and Physiology, University of North Carolina at Chapel Hill, Chapel Hill, North Carolina, United States of America; Lineberger Comprehensive Cancer Center, University of North Carolina at Chapel Hill, Chapel Hill, Chapel Hill, North Carolina, United States of America

**Keywords:** shRNA design, RNAi, Dicer, Ago2

## Abstract

Gene silencing via short hairpin mediated RNAi (shRNA) is a valuable experimental tool and has promise as a therapeutic strategy. Several shRNA platforms make use of the loop and flanking sequences from the endogenous microRNA (miRNAs) miR-30a or other miRNAs to provide an RNA structure for efficient and accurate biogenesis of the RNA trigger. However, the stem regions of these shRNAs are typically designed as perfect duplex structures which is an uncommon feature for endogenous miRNA precursors. A limitation of these designs is that shRNAs with perfect duplex stems undergo extensive stem cleavage analogous to the Dicer independent miRNA miR-451, destroying the shRNA trigger sequence that is present in the 3P arm. We employed an unbiased screen of > 9000 shRNA structures to identify features that prevent stem cleavage and promote canonical biogenesis and loading into the effector complex RISC. We find that a central stem bulge or kink reduces central stem cleavage and improves accuracy of Dicer processing. Furthermore, 9 - 10 GC nucleotides in the guide strand improves shRNA efficiency. These design rules enable more effective shRNA tools and are compatible with existing sets of optimized guide/target sequences.

## Introduction

RNA interference (RNAi) is a widely used technology to study gene function. Broadly, RNAi can be divided into two experimental approaches based on the form of the gene silencing trigger. siRNAs are short duplex RNAs that are transiently transfected into the experimental cell line. After incorporation into the RNA induced silencing complex (RISC), the guide strand of the siRNA directs RISC to complementary target RNAs and promotes silencing of the target gene. This form of RNAi is relatively short lived due to the transient nature of the siRNA trigger. In contrast, plasmid encoded shRNA triggers are capable of long term gene silencing. shRNAs are stem-loop RNAs that require cellular biogenesis pathways prior to RISC loading. shRNA expression systems based on retroviral vectors are especially valuable since they can be used for silencing gene expression long-term in cells refractory to transient transfection.

shRNA expression vectors are based on two general designs. The original shRNA vectors used the U6 or H1 promoter to drive expression of a stem loop RNA with a 19 or 29 nucleotide stem and a small loop (Brummelkamp et al. 2002; Paddison et al. 2002). These RNAs resemble miRNA precursors and are directly processed by the miRNA biogenesis enzyme Dicer prior to RISC loading. Alternative vector designs have the shRNA trigger embedded in the loop and flanking sequences of a miRNA, commonly miR-30a (shRNAmir)(Zeng et al. 2002; Silva et al. 2005). This shRNA transcript resembles a miRNA primary transcript and requires Drosha and Dicer processing prior to RISC loading. shRNAmirs are typically expressed from a RNA polymerase II promoter and therefore allow the use of the full complement of regulatable mammalian promoter systems. Additionally, the lower transcription rate of RNA polymerase II promoters and more natural loop and flanking regions result in reduced toxicity of the RNA compared to U6 based shRNA vector systems (Grimm et al. 2006; Castanotto et al. 2007; McBride et al. 2008; Shimizu et al. 2009; Ahn et al. 2011; Sun et al. 2013). Notably, both vector designs frequently place the shRNA trigger sequence on the bottom of the stem loop (3P strand) for more accurate biogenesis cleavage products. Furthermore, both shRNA designs commonly maintain perfectly duplexed stem structure unlike endogenous miRNAs (including miR-30a). While retroviral shRNA systems have found widespread use in biomedical research they often display incomplete target gene knockdown, especially when integrated at a single genomic copy (Bhinder and Djaballah 2013; Bhinder et al. 2014). Several strategies have been developed to identify strong target sites within mRNAs, though the shRNA itself retained the basic structure (Fellmann et al. 2011; Premsrirut et al. 2011; Fellmann et al. 2013).

We have previously developed next generation sequencing technologies for the study of miRNA biogenesis (Newman et al. 2011). Here, we applied these approaches to the analysis of shRNA biogenesis. We find that the vast majority of shRNA precursors are internally cut on the 3P stem analogous to the Dicer-independent miRNA miR-451. Since shRNA vector systems are designed with the gene silencing trigger on the 3P strand, this will limit the amount of trigger RNA that can be loaded into RISC. To improve shRNA biogenesis efficiency we developed an unbiased functional screen to identify structural features that promote biogenesis and RISC loading. We report that central mismatches are essential to prevent internal 3P cleavage and that additional features including flanking mismatches and GC content improve shRNA efficacy.

## Results

We investigated the production of mature shRNA triggers from the widely used pTRIPZ retroviral vector. This vector has the shRNA duplex sequence within the context of the miR-30a loop and flanking sequences. Transcription is driven from a tetracycline inducible promoter. We studied two moderately weak shRNAs that target the Col1A1 and Fox3 mRNAs. Single copy integration into NIH-3T3 cells led to 51 and 43% knockdown, respectively, for the two shRNAs (Figure 1A). As expected, introduction of additional copies of the shRNA vector by transient transfection improved target gene knockdown. We prepared small RNA libraries for next generation sequencing to quantify the amount of mature shRNA trigger present in stably expressing cells. We observed 1720 and 2578 reads per million for the Col1A1 and Fox3 shRNAs, respectively, in cells with single copy integration of the corresponding TRIPZ vector. This is in contrast to endogenous miRNAs that in some cases had 100 fold more mature sequence reads. Notably, miR-21a is a miRNA present at a single genomic locus yet had greater than 200,000 normalized sequence reads.

**Fig. 1.**
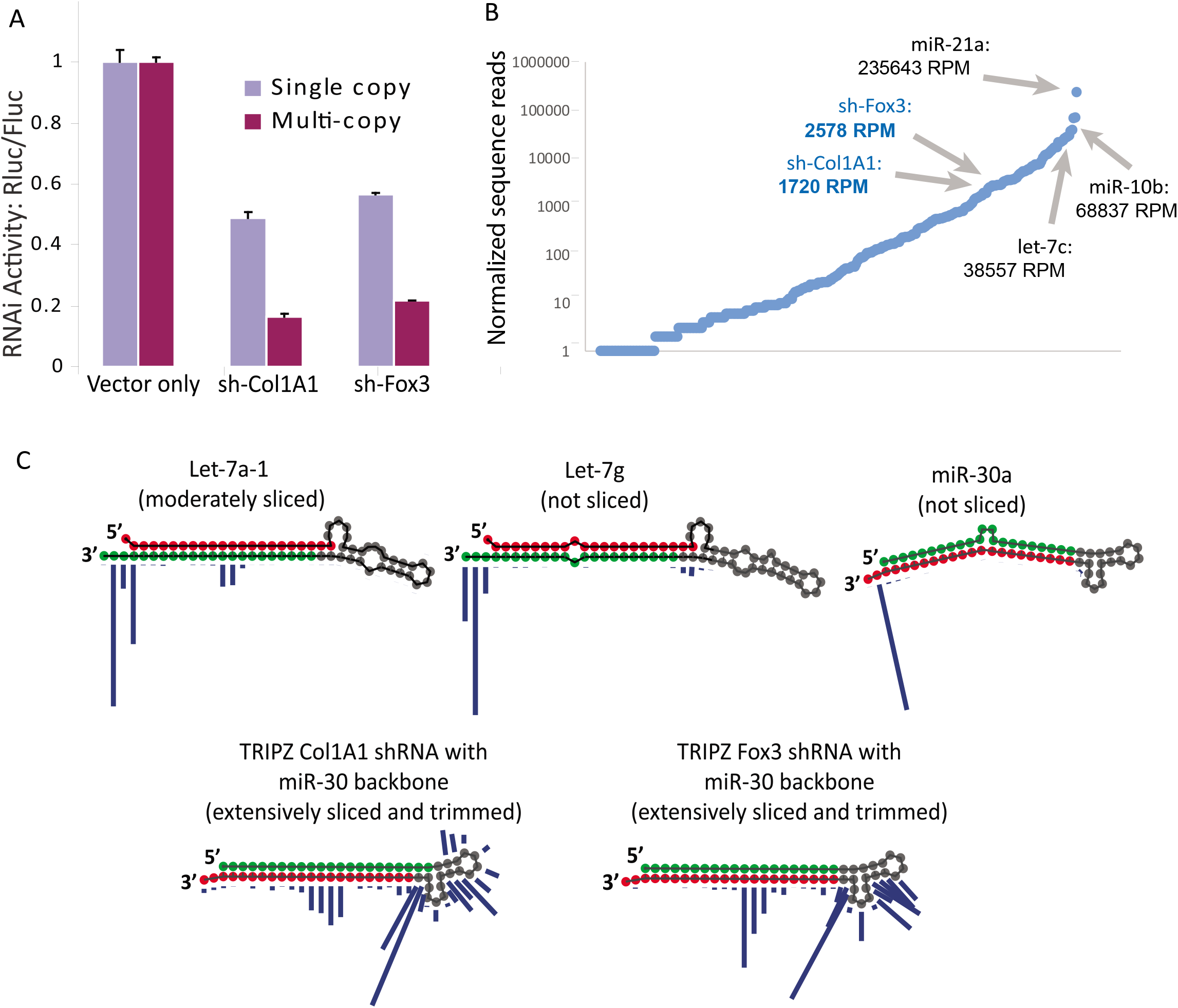
Stem cleavage reduces biogenesis efficiency of shRNAs. **(A)** Knockdown activity for two different shRNAs was determined using a luciferase reporter vector. NIH-3T3 cells were stably transduced with shRNA retrovirus at single copy integration and target knockdown measured. Multi-copy knockdown was determined by transient transfection of additional shRNA vector into cells. **(B)** Small RNAs from shRNA transduced cells were sequenced using the Illumina platform. Normalized miRNA and shRNA reads are plotted in rank order. **(C)** Precursor miRNAs and shRNAs were sequenced using a customized Illumina protocol. Read counts for individual miRNAs/shRNAs were counted for each terminal nucleotide position on the precursor sequence. The height of each blue bar indicates the relative read count for precursors that terminate at the indicated 3’ position. Three representative miRNAs are shown along with the shRNAs.

We next sought to identify causes for the low yield of mature shRNA trigger in virus transduced cells. We employed our previously developed deep sequencing approach specific for miRNA precursors (Newman et al. 2011). This approach provides not only read counts for specific precursor sequences but also provides information about internal 3P stem cleavage, trimming, and nontemplated nucleotide addition. For example, analysis of precursor reads for the miRNA Let-7g demonstrates that almost all precursors are full length sequences, with < 1 percent of the sequences trimmed and/or cut internally in the 3P stem or near the loop (Figure 1C, blue bars indicate relative reads at each nucleotide position). Let-7a1 has a greater fraction of precursors that are cleaved internally; however, most precursors remain full length. The increase in internal cleavage for Let-7a1 is likely due to the perfect duplex structure of the stemloop, a feature known to promote Ago2 mediated cleavage.

We then analyzed precursor reads for the two shRNAs. Over 99 percent of reads had evidence of internal cleavage and trimming. This cleavage pattern is reminiscent of the Dicer independent miR-451 biogenesis and may be related to a reported Ago2-based precursor surveillance system (Diederichs and Haber 2007; Cheloufi et al. 2010; Cifuentes et al. 2010; Yang and Lai 2010; Liu et al. 2012; Liu et al. 2014). Notably, the endogenous precursor for miR-30a, on which the TRIPZ system is based, contains an asymmetric bulge at nucleotides 12/13 of the 5P arm and is resistant to internal cleavage. However, when the miR-30a precursor was adapted to the shRNA vector system the central stem bulge was not retained; only the loop and flanking sequences were transitioned to pTRIPZ. The stem region of these shRNAs are perfect duplexes and therefore can be substrates of internal Ago2 cleavage. This is a design limitation since the trigger strand is the 3P strand of the stem. Any internal cleavage will destroy the trigger, reducing potency and potentially increasing incorporation of the 5P strand with concomitant off-target effects. It should be noted that trigger strand cleavage is not only observed with TRIPZ. We find extensive stem cleavage with the U6 based pLKO vector - which also has a perfect duplex stem (data not shown).

To directly test whether the perfect duplex stem was promoting internal cleavage we modified the shRNA vector to include a central bulge in the duplex region in the range of positions 9-11, positions that inhibit Ago2 cleavage of siRNA passenger strands, target mRNAs, and miR-451 biogenesis (Martinez et al. 2002; Leuschner et al. 2006; Cifuentes et al. 2010; Yang and Lai 2010). We introduced these mismatches in the 5P non-trigger (passenger) strand of both the Col1A1 and Fox3 shRNA vectors. The 3P guide sequence was unchanged to allow direct comparison to the original structure. Precursor sequencing of these modified shRNAs demonstrated that > 99 percent of reads were full length with very limited internal cleavage (Figure 2A). We directly compared knockdown efficiency of the stem-bulged shRNAs with perfect duplex shRNAs. At single copy integration, bulged shRNAs led to increased knockdown potency (Figure 2B).

**Fig. 2.**
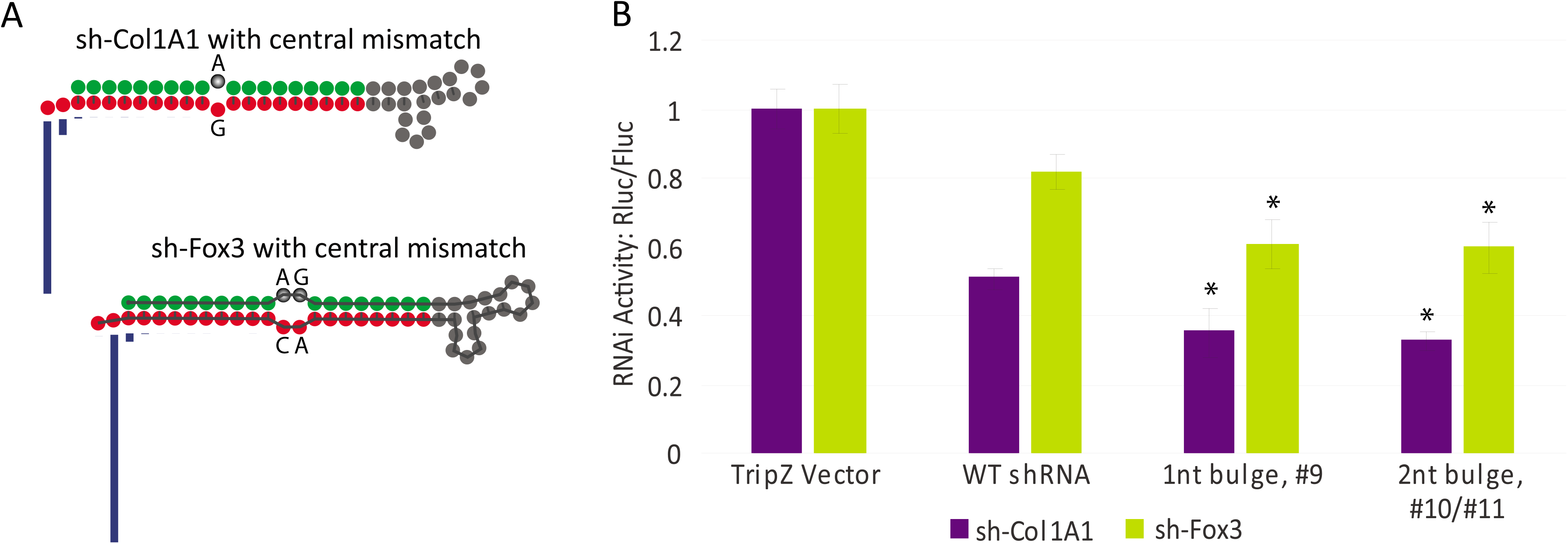
shRNA stem-central bulges prevent internal cleavage and promote knockdown activity. **(A)** Mutations were introduced into precursor sequences to create a central bulge in the stem loop structure. A single-nucleotide bulge at #9, counting from the 5’ terminal end, and two nucleotide bulges at #10/11 were introduced in the 5P arm to allow identical trigger sequences. Precursor shRNAs were sequenced and read counts for individual shRNAs were counted for each terminal nucleotide position on the precursor sequence. The height of each blue bar indicates the relative read count for precursors that terminate at the indicated 3’ position. **(B)** Knockdown activity was determined of mutant shRNAs structures from (A). shRNAs were transduced at single copy and reporter activity measured.

While these rational design changes improved shRNA performance, we wanted to identify other structural features that would further optimize shRNA biogenesis and RISC loading. Previous studies have employed screening of libraries of miRNA sequences to identify features that improved Drosha and Dicer cleavage. This has led to the identification of primary sequence motifs including CNNC, GHG, and GUG, present in flanking regions and loops, respectively, that improve Drosha binding and cleavage (Auyeung et al. 2013; Fang and Bartel 2015; Kwon et al. 2019). We wanted to identify features that improve the entire biogenesis pathway, and avoid Ago2 slicing of the stem, to maximize target knockdown potential. We devised an unbiased functional screen to identify these optimal shRNA structures, summarized in Figure 3. Through the use of degenerate nucleotides, we generated a library of different shRNA structures within the TRIPZ vector. These structures included central bulge options at stem positions 9-10, asymmetrical mismatches that lead to kinked stem structures, and mismatches or G·U wobble pairing in 5’ and 3’ stem regions (nucleotides 3,4,5 and 14,15,16, counting from the 5’ precursor phosphate). Libraries were based on shRNAs against three target proteins, Gef7, Tiam1, and WasL. These parent shRNAs ranged from high to low in thermodynamic pairing energy, to reveal differential effects on mismatches. Low pairing energy was maintained proximal to the loop, to drive the desired 3P strand selection into Ago. All degenerate nucleotides were introduced on the 3P trigger strand; the 5P strand was unaltered for all structures (Fig 3B). This allows identification of the preferred variants by sequence analysis of mature shRNAs generated from the 3P strand. We introduced these libraries of over 9000 possible structural variants into NIH-3T3 cells by single-copy retroviral transduction. To identify structural variants that enhanced biogenesis and RISC loading, we isolated Ago2 associated RNAs by immunoprecipitation, and identified enriched degenerate nucleotide structures, normalized to the sequenced provirus population.

**Fig. 3.**
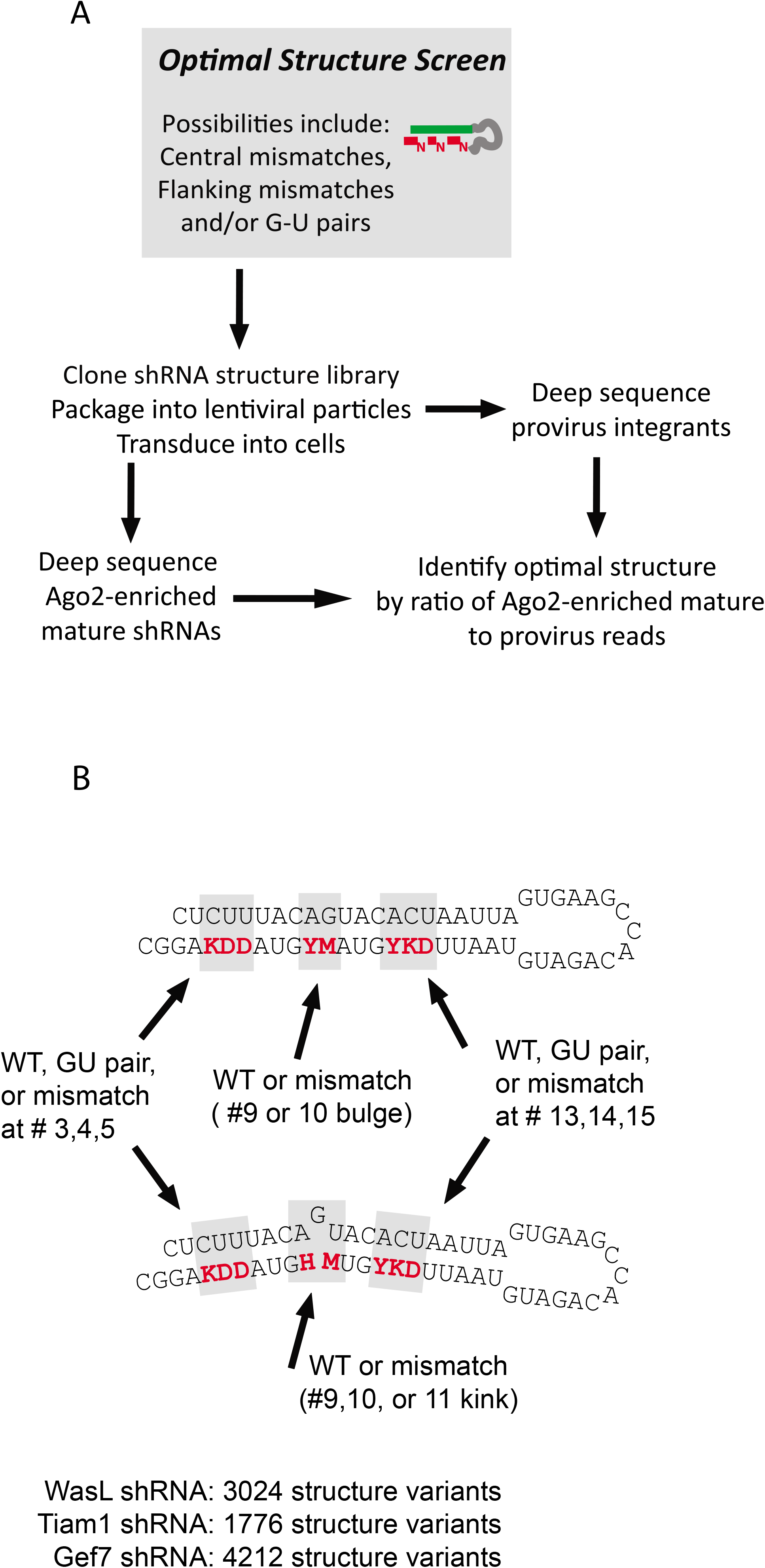
Unbiased screen design for optimized shRNA structures. **(A)** The flow chart for our shRNA structure screen is shown. shRNA structure libraries containing central bulges or kinks, and additional flanking bulges were generated using degenerate oligonucleotides. Libraries were constructed in the miR-30 TRIPZ vector backbone. Structure libraries were packaged into virus and transduced into NIH-3T3 cells. Ago2 associated small RNAs were isolated and identified by deep sequencing. Optimal structures were identified by normalizing deep sequencing reads of Ago-associated guide strands to provirus reads. **(B)** The structural variants in the library are illustrated with degenerate positions in red, for a kinked and non-kinked structure. Base position numbering starts at 1 for the 5P precursor end. Position numbers refer to 3P bases paired to the indicated number on the 5P strand. Oligonucleotide sequences are listed in the Supplement.

Features that lead to enrichment of shRNAs are shown in Figure 4. Each shRNA library for Gef7, Tiam1, and WasL, are shown as separate heatmaps. Vertical columns represent the possible central bulges or kinks, and each row represents possible mismatches that flank the central bulge. Up to six flanking mismatches are possible, for 64 total combinations, for each central bulge/kink sup-population. For this analysis, G·U wobble and A·U pairing were considered equivalent. Therefore, most central/flanking mismatch combinations had up to eight variants, varying by A·U or G·U at non-mismatched positions. This allowed averaging of variants with the same mismatches using linear regression, improving the robustness of the analysis (see Supplementary Figure 1). As shown in parent sequences at the bottom of the Figure, the Gef7 shRNA parent has the highest energy pairing in the stem (G·C pairs shown in gray boxes) and the WasL has the lowest. This feature drove the preference for flanking mismatches, and to a lesser extent, central bulge/kink structures. The low energy stem of WasL was not tolerant of central bulges or kinks, but one or two flanking mismatches lead to efficient processing. The fully wild type sequence, however, was still highly favored. For Tiam1, one or two flanking mismatches, with or without a bulge at position 9, were most efficient. The high GC content of the Gef7 stem was permissive of more mismatches. In fact, the fully wild type (parent) sequence was not efficiently processed. The preferred structure had four flanking mismatches along with no central bulge, or a single nucleotide bulge at 9 or 10. Interesting, kinked structures were also enriched. The TRIPZ vector was designed around the miR-30a backbone, and this miRNA has a two nucleotide kink in the stem. Kinked structures were not tolerated in the Tiam1 or WasL shRNA libraries, presumably due to further stem destabilization due to higher AT content. Specifically, the Gef7 9,10 kink has a G·C pair on each side of the kink that might maintain the duplex structure. miR-30a also has this flanking GC feature, with 8 total G·C pairs in the precursor stem. miR-30a predominantly loads the 5P stem into Ago, however, so the natural miRNA may not fully translate to the shRNA structure.

**Fig. 4.**
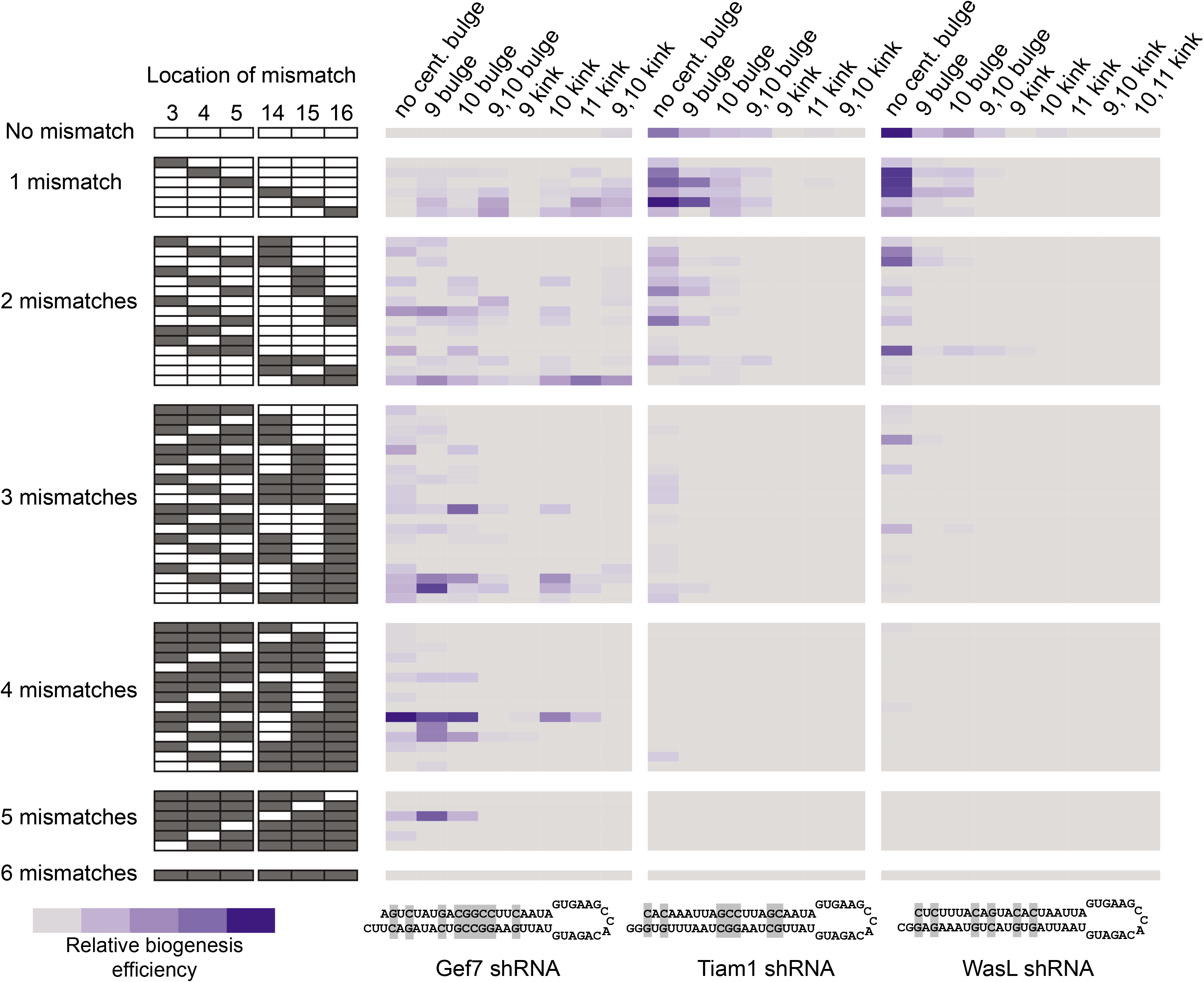
Central bulges or kinks can improve efficiency of shRNA structural variants. Relative enrichment of Ago2 loaded shRNA variants is shown in a heatmap. Libraries were based on Gef7, Tiam1, and WasL parent shRNAs and are shown on separate heatmaps. Vertical columns represent different central bulge/kink structures, and rows represent the presence of mismatches flanking the central structural motif. The location of flanking mismatches is shown at the far left black/white heatmap, with back representing a mismatch. Parent shRNA sequences are shown at bottom, with GC pairing in the stem duplex highlighted.

Further breakdown of the data at each flanking variant position for G·U pairing revealed interesting trends. As shown in Figure 5, mismatches at all six positions were prevalent for Gef7, essentially recapitulating the heatmap in Figure 4. More interesting, however, was the strong preference for G·U wobble at two and three positions for Tiam1 and WasL, respectively. In all cases except one, a U·A pair (WT) is replaced with U·G. This would have a limited effect on the thermodynamics of the stem duplex, but would increase GC content of the mature shRNA trigger. Figure 5B shows the enrichment of shRNA variants based on GC content of the mature shRNA. Increases in GC content, either by replacing a A·U with a G·U pair, or an A·U to a A·C mismatch, are counted as gains in GC content, while a G·C to G·A switch is counted as a loss. Specifically, we are analyzing the effects that are driven by GC gains in the mature guide RNA, but will not lead to increased GC pairing in the duplex stem. The WasL shRNA, with a lower GC content in the parent sequence, showed enrichment for sequences with a GC gain of 3-4 nucleotides. In contrast, the higher GC Gef7 shRNA does not enrich for a gain. These data support a preference for a guide RNA GC content of ~9, regardless of the pairing energy of the duplex stem.

**Fig 5.**
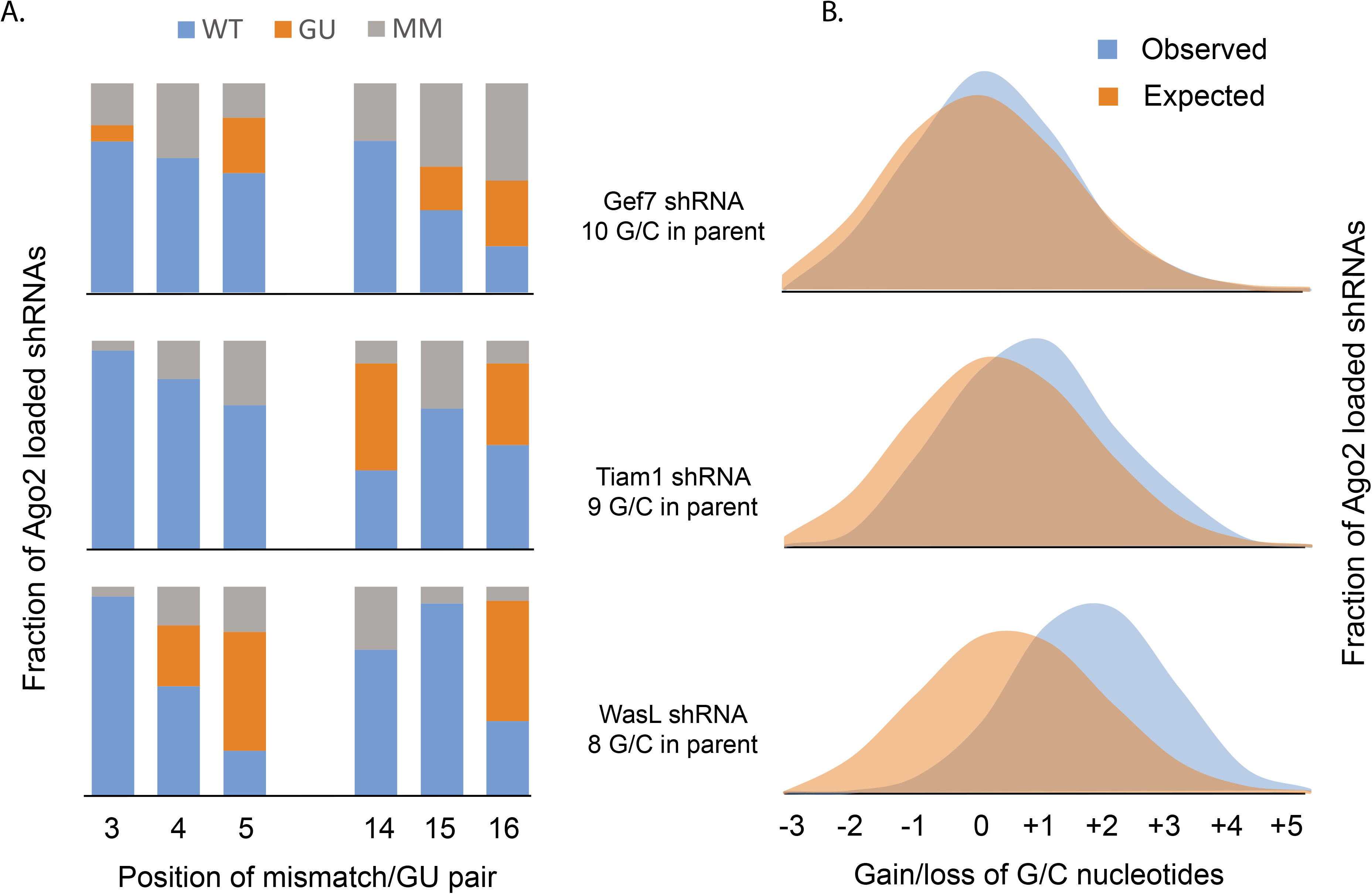
Increased GC content and G·U wobbles improve shRNA efficiency. **(A)**. Relative fraction of enriched shRNA structures with G·U wobble, mismatch, or wild-type (parental) sequence at indicated position. Asterisk indicates the position where the G·U has a significant impact on stem pairing energy (G·C to G·U transition), all other positions have A·U to G·U transitions. **(B)** Relative fraction of enriched shRNA structures relative to gain or loss of GC content in the mature shRNA. All variants that increased GC content in the stem, including A·U to G·U transitions and mismatches that introduce a C in the 3P strand, were counted as gains of GC content, independent of changes in stem base pairing energy. Expected fraction is based on total number of variants at each position.

Another consideration for shRNA design is the accuracy of cleavage. Off-target effects of shRNAs can be mediated by a miRNA-like mechanism, where the “seed” sequence is the primary determinant of targeting. Due to this effect, heterogeneity of the 5’ end of the mature guide RNA will lead to complex and unpredictable off-target silencing. We analyzed the accuracy of the 5’ cleavage event in Ago2 loaded shRNA variants, binned according to central bulge/kink structure. As shown in Figure 6, perfect duplex shRNA structures had a mixture of cleavage sites, depending on sequence. The WasL shRNA had good accuracy for most central structures, even without the optimal U in position 1 of the mature shRNA. The Gef7 and Tiam1 shRNAs were cleaved at two or more sites for many central structures. These shRNAs should have a U at position 1 if processed as expected, thus the secondary cleavage sites are unexpected. Kinked structures were more accurately processed for all shRNA libraries, however this occurs with a loss of efficiency for low GC stem shRNAs (WasL and Tiam, Figure 4).

**Fig 6.**
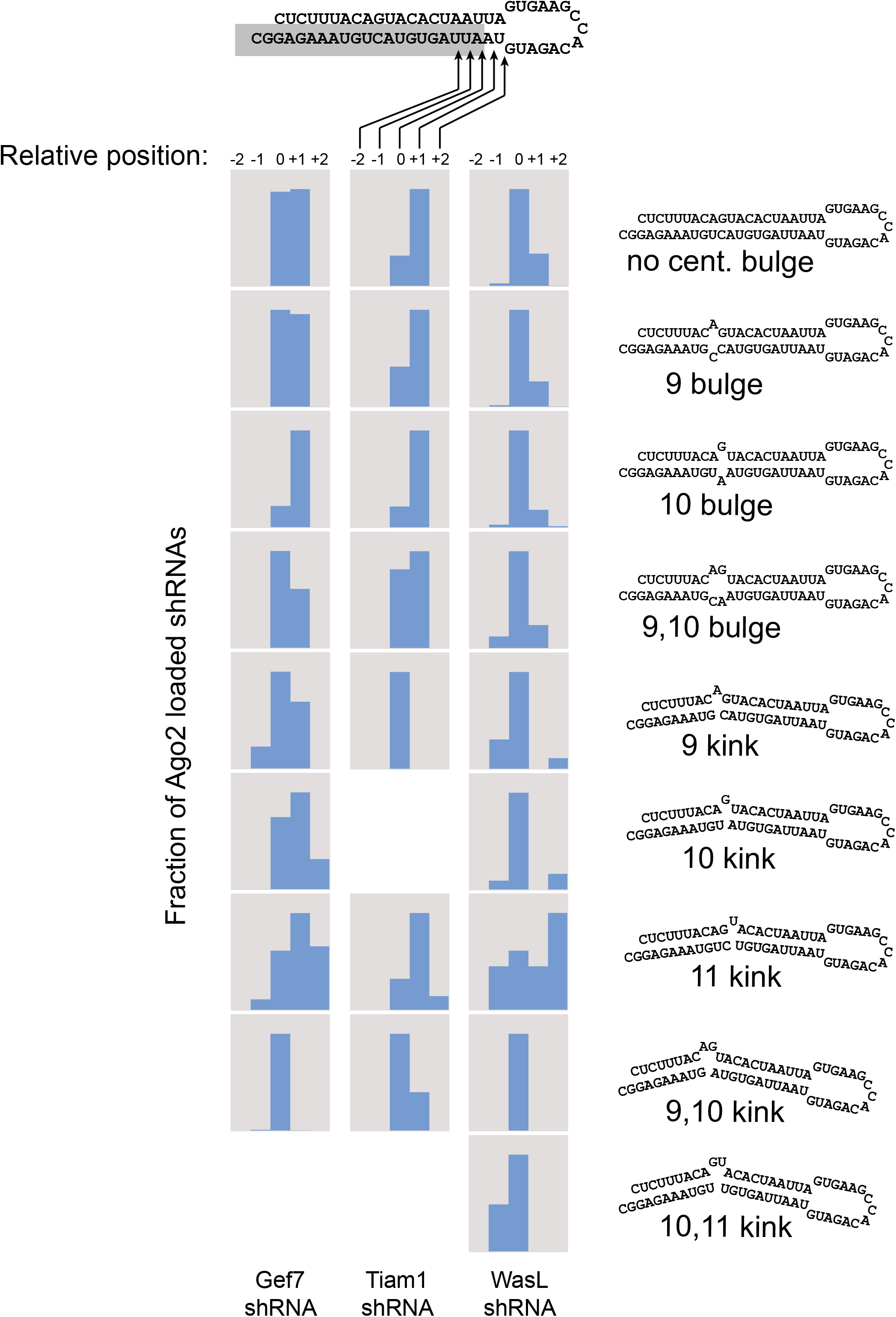
Central structural motifs dictate 5’ cleavage accuracy. Fraction of enriched variants are shown in the blue bar chart at each cleavage position. The parent shRNA for WasL is shown for reference, with the gray shaded region the desired mature shRNA. Each type of central bulge/kink is shown in a separate row, as labeled on the right. Sequences for WasL structures are shown at right for illustrative purposes, Gef7 and Tiam1 are similar.

## Discussion

In this report we describe a functional screen for the optimization of shRNA structures. Our approach was designed to interrogate all steps in shRNA processing and Ago2 loading, maximizing the potential for target gene knockdown. We demonstrate that Ago2-mediated cleavage of perfect duplex stem-loops limits the yield of functional shRNA guide RNAs. Central bulges or kinks in the stem can prevent Ago2-mediated internal cleavage and trimming of shRNA precursors, as has been previously described in approaches to investigate Dicer activity or improve shRNA function (Vermeulen et al. 2005; Vlassov et al. 2007; Boudreau et al. 2008; Wang et al. 2008; Schopman et al. 2010; Wu et al. 2011; Jensen et al. 2012; Matveeva et al. 2012; Denise et al. 2014; Fowler et al. 2016; Watanabe et al. 2016; Xie et al. 2020). shRNA stem bulges have also been hypothesized to decrease thermodynamic pairing energy. G·U wobbles have been described in stems to promote Dicer processing, may have the same effect as bulges when it comes to guide/passenger strand selection, and have been reported to inhibit Dicer independent AgoshRNA activity (Harborth et al. 2003; Matranga et al. 2005; Okamura et al. 2009; Gu et al. 2011; Gu et al. 2014; Ma et al. 2014; Kampmann et al. 2015; Liu et al. 2015). Our screening approach allows evaluation of GC content of the stem and its impact on biogenesis efficiency and accuracy on central bulge/kink structures. Low GC content disfavors a central bulge or kink, but that factor needs to be weighed against cleavage accuracy and resultant potential for off-target effects. Interestingly, asymmetric mutations in the stem that leads to a kinked structure has previously been reported to reduce accuracy of Dicer cleavage; however our data suggests that this is dependent on kink position and stem GC content (Starega-Roslan et al. 2011). Interestingly, the Ago-2 slicing pathway has been successfully harnessed for use via Dicer-independent AgoshRNAs (Herrera-Carrillo et al. 2014; Harwig et al. 2015; Kaadt et al. 2019). Ago2 cleavage of precursors is dependent on a perfect duplex stem, a feature that is absent in most endogenous miRNA precursors. In the case of miR-451, Ago2 cleavage of the 3P stem can lead to productive mature miRNA, since the mature strand is on the 5P stem. The most common shRNA vector designs, however, place the active shRNA trigger on the 3P stem. Thus, Ago2 cleavage would destroy the trigger and compromise the potency of the shRNA. While the structural features that promote Ago2 cleavage of precursors has been known, our deep sequencing results described herein demonstrate the extent of precursor loss and reduces the efficiency of many conventional shRNA designs.

Our functional screen also demonstrated that biogenesis and/or Ago loading are dependent on guide strand GC content. This effect is independent of GC content in the duplex stem itself, therefore would not be expected to impact passenger strand unwinding. As our screen data shows, GC content in the guide can be increased, without increasing stem GC pairing, by conversion of UA pairs to UG (A to G transition on guide strand, across from U). Interestingly, large-scale identification of effective shRNA guide sequences has revealed highly AT rich target sequences (Knott et al. 2014; Pelossof et al. 2017). The resulting guide RNAs can be adjusted for increased GC content, using A to G transitions, without changing the target site. This allows our design optimizations to be readily adapted to existing sets of guide RNAs.

While some of our reported improvements have been previously demonstrated, the adoption of these design changes to commercial shRNA vector systems is still underway. The miR-30-based TRIPZ vector as well as U6 based vectors (pLKO from the RNAi Consortium) are still designed with perfect duplex stems. One reason for the resistance to new structures is that shRNA libraries would need to be redeveloped. Furthermore, it is not clear that previous studies were able to maximize the design improvement due to the rational method of design changes. The unbiased structure library screen that we report here is an alternative strategy to develop an optimized structure based on RISC loading enrichment.

While our approach has the advantage that it interrogates all biogenesis steps in a single experiment, it has limitations. Since the shRNAs are integrated at single-copy in the mouse genome, the majority of Ago2-associated RNAs will be endogenous miRNAs. Thus, the approach cannot reveal quantitative effects in weakly processed structures, but will highlight sequence and structural features that improve biogenesis efficiency. However, the overall goal of our study is to model single-copy shRNA potency, and high copy shRNA expression might favor different structural features. Previous studies have used libraries of artificial miRNA sequences to probe Drosha and Dicer processing motifs (Auyeung et al. 2013; Fang and Bartel 2015; Kwon et al. 2019). These studies were designed to capture and identify the library RNAs without the background of endogenous miRNAs, allowing complete saturation of the screen. These studies did not interrogate the full biogenesis and RISC loading pathway, however. We anticipate that integration of our findings with these studies and target site and guide strand identification will lead to optimized designs for shRNAs.

## Materials and Methods

### Cell culture, transfection, and transduction

NIH-3T3 fibroblasts and 293T packaging cells were cultured as described by ATCC.org. To generate virus, 293T cells were triple-transfected with pTRIPZ or pLKO vector, delta NRF packaging plasmid, and VSVG envelope. NIH-3T3 cells were transduced with virus as described by Dharmacon, using 1:3 serial dilutions into 6-well or 10 cm plates to obtain singlecopy MOI. Cells were selected and maintained in 1.25 ug/ml puromycin. For reporter assay, 1 x 10^5^ transduced cells were plated in 24-well format on Day 1; shRNA expression was induced after cell attachment with 1 ug/ml doxycycline (replenished every 24 hours until lysate harvest). On Day 2, psiCHECK 2.0 constructs were transfected using Lipofectamine 2000. On Day 3, media was changed, and the lysate was harvested on Day 4 using Promega Passive Lysis Buffer. For co-transfection studies, 1ug/ml doxycycline was present during transfection of both the TRIPZ and psiCHECK 2.0 plasmids, otherwise usage was identical to Day 3 onward of the transduction protocol. All luciferase assays within a single panel of a figure compare singlecopy integrants of the same passage number (i.e. control, parental, and experimental shRNAs were transduced side by side), and exposed to doxycycline and puromycin premixed prior to use into a common source. Error bars are the standard deviation of technical replicates of three independent transfections of reporter.

### Retroviral and reporter constructs

Single shRNAs and structure libraries were cloned using a modification of a published protocol, using oligonucleotide sequences shown in Supplementary Table 1. Specifically, 97mers containing the entire stem-loop sequence were used as PCR templates, and flanking primers added EcoR1 and Xho1 sites. In the case of structure libraries, 97mers contained degenerate nucleotides at positions in the 3P stem. To generate library variants with different total length (no kink vs single nucleotide kink vs two nucleotide kink) the degenerate oligonucleotides were pooled before PCR. Products were ligated into the pTRIPZ vector (Open Biosystems), transformed, and plated. Colonies were scraped and plasmid directly purified. Sequencing of plasmid libraries demonstrated full coverage of library variants.

pKLO plasmids were obtained from the UNC Functional Genomics Core. Guide and passenger strand sequences were obtained from the RNAi Consortium, pLKO.1-based plasmids TRCN0000090506 (Col1A1) and TRCN0000254936 (Fox3). To generate matched TRIPZ constructs, guide sequences were converted to the miR-30 backbone, and 97mers cloned as described above.

For RNAi reporter vectors, perfectly complementary 1X targets were designed to anneal and produce sticky ends ready for psiCHECK 2.0 XhoI/NotI. All shRNAmir and reporter oligonucleotides used can be found in Supplementary Table 1.

### Precursor next generation sequencing

For precursor-specific NGS, RNA was isolated from transduced NIH-3T3 cells. Libraries were generated as previously described using pooled miRNA-specific 5’ oligos (Newman et al. 2011). Included were oligos directed at the shRNA 5P stem. Analysis was performed as described, mapping individual shRNA or miRNA reads against expected matches, and counting 3’ terminal end position.

### Structure library next generation sequencing and analysis

NIH-3T3 cells transduced with shRNA structure libraries were induced with doxycycline as described above. Cells were lysed in buffer A: 50 mM Hepes pH 7.5, 100 mM NaCl, 0.5% NP40. Ago-associated RNAs were immunopurified using Protein A Dynabeads (Life Technologies) and the 2A8 antibody, preferential for Ago2 (Diagenode) in Buffer A. Beads were washed 4x in Buffer A and 2x in Buffer A supplemented adjusted to 500 mM NaCl. RNA was extracted using Trizol (Invitrogen). Small RNA NGS libraries originated from 5.0 ug total RNA using a method similar to that previously published and similar to the Illumina protocol (Pfeffer et al. 2005). Briefly, RNA adapters were ligated to the 5’ and 3’ ends of RNA targets, followed by reverse transcription and PCR. Libraries were gel purified and sequenced on a HiSeq 2000 (Illumina). Primers that amplified the flanking regions of shRNAs were used to verify plasmid library screen depth, as well as to identify and quantify genomic integrants in the population for normalization; see S1 Table.

Raw sequence reads were processed using a custom bioinformatics pipeline, as previously published (Newman et al. 2011). Mature shRNAs from the library screen were mapped to the originating structural variant. Reads for each variant were normalized to the amount of integrated provirus for that variant using regression analysis. Reads were further binned by 5’ cleavage location and/or stem GC content, as depicted in the corresponding figures.

## Acknowledgements

Work in this study was funded by the NIH U01-HL111018 and IMAT program R21-CA196379.

## Supporting Information

**S1 Fig.**
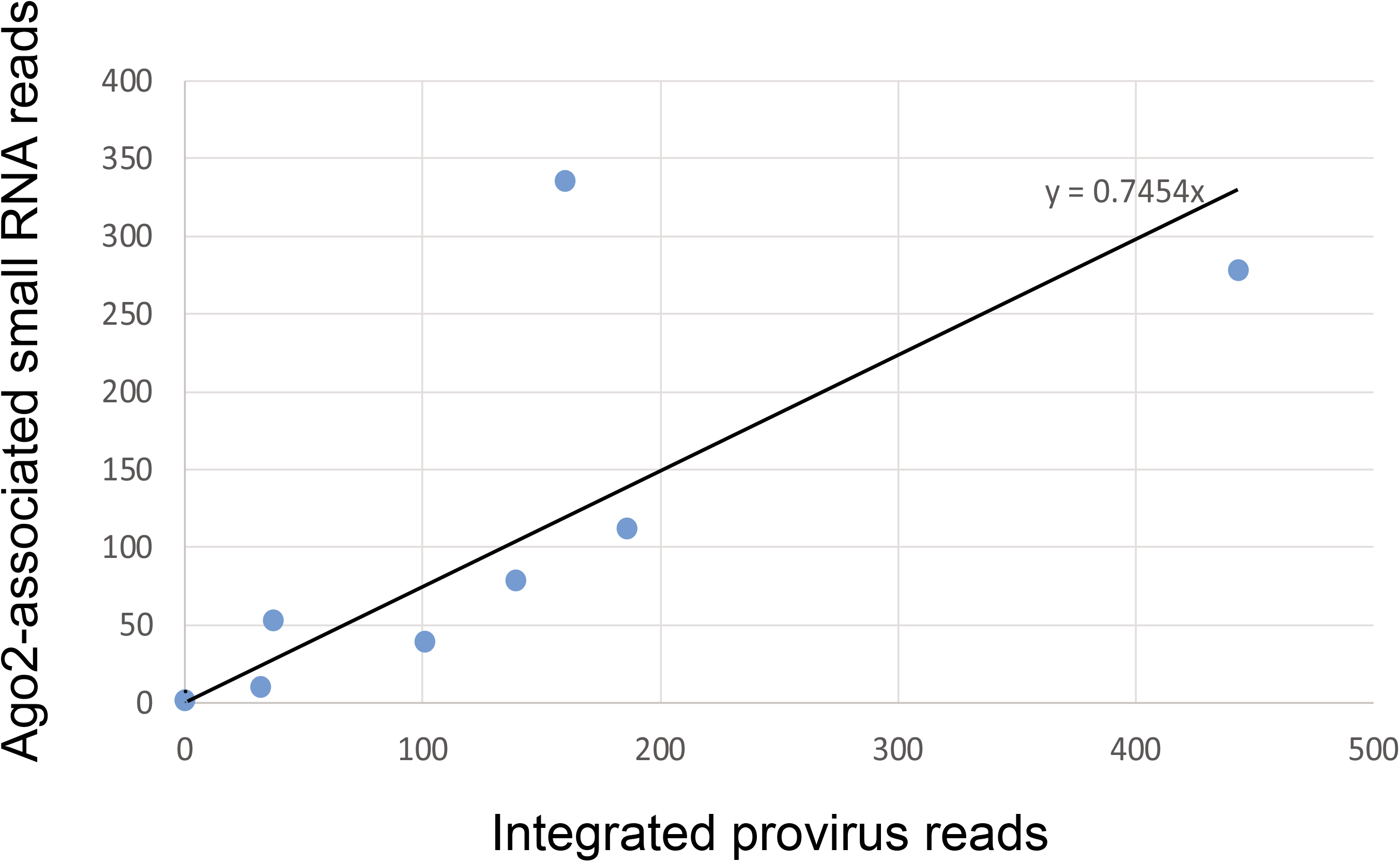
Least squares fit for enrichment values. **(A)** shRNA variants for WasL library, with no central bulge/kink and no flanking mismatches, are plotted as Ago2 small RNA reads versus integrated virus normalizer reads. Eight variants are in the library, varying only by the presence of A·U to G·U transitions at three positions. Linear regression line is shown, with slope value.

**S1 Table.**
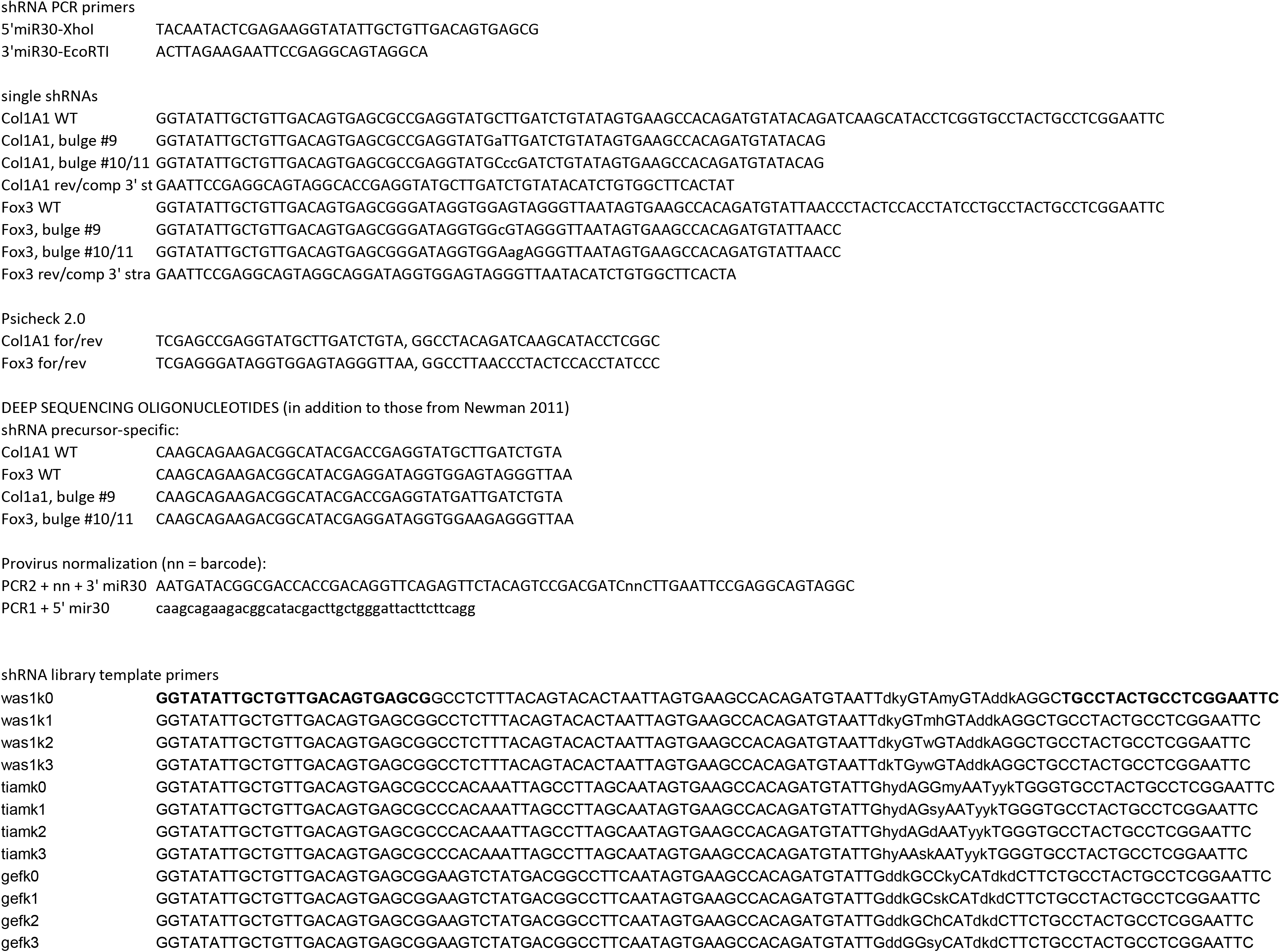
Oligonucleotides used to produce constructs and deep sequencing libraries. Amplification primers for shRNA libraries are shown. Primers were used as templates in a PCR reaction to introduce restriction-enzyme substrates for subsequent steps. PCR products were subcloned into TRIPZ. For specific shRNAs, in some cases the template primers were in two sections, using overlap-extension to generate final template. For cloning of Col1a1 and Fox3 reporters, psiCHECK 2.0 primers were annealed, diluted for a 3:1 insert:vector ratio and ligated into vector. Pre-seq specific primers were used to select for shRNA precursors, and were added to a multiplexed set of primers as previously published (Newman et al. 2011). Provirus normalization primers were used to amplify genomic DNA for subsequent enrichment studies vs. Ago-associated triggers.

## References

Ahn M, Witting SR, Ruiz R, Saxena R, Morral N. 2011. Constitutive expression of short hairpin RNA in vivo triggers buildup of mature hairpin molecules. Hum Gene Ther 22: 1483–1497.

Auyeung VC, Ulitsky I, McGeary SE, Bartel DP. 2013. Beyond secondary structure: primary-sequence determinants license pri-miRNA hairpins for processing. Cell 152: 844–858.

Bhinder B, Djaballah H. 2013. Systematic analysis of RNAi reports identifies dismal commonality at gene-level and reveals an unprecedented enrichment in pooled shRNA screens. Comb Chem High Throughput Screen 16: 665–681.

Bhinder B, Shum D, Djaballah H. 2014. Comparative analysis of RNAi screening technologies at genome-scale reveals an inherent processing inefficiency of the plasmid-based shRNA hairpin. Comb Chem High Throughput Screen 17: 98–113.

Boudreau RL, Monteys AM, Davidson BL. 2008. Minimizing variables among hairpin-based RNAi vectors reveals the potency of shRNAs. RNA 14: 1834–1844.

Brummelkamp TR, Bernards R, Agami R. 2002. A system for stable expression of short interfering RNAs in mammalian cells. Science 296: 550–553.

Castanotto D, Sakurai K, Lingeman R, Li H, Shively L, Aagaard L, Soifer H, Gatignol A, Riggs A, Rossi JJ. 2007. Combinatorial delivery of small interfering RNAs reduces RNAi efficacy by selective incorporation into RISC. Nucleic Acids Res 35: 5154–5164.

Cheloufi S, Dos Santos CO, Chong MM, Hannon GJ. 2010. A dicer-independent miRNA biogenesis pathway that requires Ago catalysis. Nature 465: 584–589.

Cifuentes D, Xue H, Taylor DW, Patnode H, Mishima Y, Cheloufi S, Ma E, Mane S, Hannon GJ, Lawson ND et al. 2010. A novel miRNA processing pathway independent of Dicer requires Argonaute2 catalytic activity. Science 328: 1694–1698.

Denise H, Moschos SA, Sidders B, Burden F, Perkins H, Carter N, Stroud T, Kennedy M, Fancy SA, Lapthorn C et al. 2014. Deep Sequencing Insights in Therapeutic shRNA Processing and siRNA Target Cleavage Precision. Mol Ther Nucleic Acids 3: e145.

Diederichs S, Haber DA. 2007. Dual role for argonautes in microRNA processing and posttranscriptional regulation of microRNA expression. Cell 131: 1097–1108.

Fang W, Bartel DP. 2015. The Menu of Features that Define Primary MicroRNAs and Enable De Novo Design of MicroRNA Genes. Mol Cell 60: 131–145.

Fellmann C, Hoffmann T, Sridhar V, Hopfgartner B, Muhar M, Roth M, Lai DY, Barbosa IA, Kwon JS, Guan Y et al. 2013. An optimized microRNA backbone for effective single-copy RNAi. Cell Rep 5: 1704–1713.

Fellmann C, Zuber J, McJunkin K, Chang K, Malone CD, Dickins RA, Xu Q, Hengartner MO, Elledge SJ, Hannon GJ et al. 2011. Functional identification of optimized RNAi triggers using a massively parallel sensor assay. Mol Cell 41: 733–746.

Fowler DK, Williams C, Gerritsen AT, Washbourne P. 2016. Improved knockdown from artificial microRNAs in an enhanced miR-155 backbone: a designer’s guide to potent multi-target RNAi. Nucleic Acids Res 44: e48.

Grimm D, Streetz KL, Jopling CL, Storm TA, Pandey K, Davis CR, Marion P, Salazar F, Kay MA. 2006. Fatality in mice due to oversaturation of cellular microRNA/short hairpin RNA pathways. Nature 441: 537–541.

Gu S, Jin L, Zhang F, Huang Y, Grimm D, Rossi JJ, Kay MA. 2011. Thermodynamic stability of small hairpin RNAs highly influences the loading process of different mammalian Argonautes. Proc Natl Acad Sci U S A 108: 9208–9213.

Gu S, Zhang Y, Jin L, Huang Y, Zhang F, Bassik MC, Kampmann M, Kay MA. 2014. Weak base pairing in both seed and 3’ regions reduces RNAi off-targets and enhances si/shRNA designs. Nucleic Acids Res 42: 12169–12176.

Harborth J, Elbashir SM, Vandenburgh K, Manninga H, Scaringe SA, Weber K, Tuschl T. 2003. Sequence, chemical, and structural variation of small interfering RNAs and short hairpin RNAs and the effect on mammalian gene silencing. Antisense Nucleic Acid Drug Dev 13: 83–105.

Harwig A, Herrera-Carrillo E, Jongejan A, van Kampen AH, Berkhout B. 2015. Deep Sequence Analysis of AgoshRNA Processing Reveals 3’ A Addition and Trimming. Mol Ther Nucleic Acids 4: e247.

Herrera-Carrillo E, Harwig A, Liu YP, Berkhout B. 2014. Probing the shRNA characteristics that hinder Dicer recognition and consequently allow Ago-mediated processing and AgoshRNA activity. RNA 20: 1410–1418.

Jensen SM, Schmitz A, Pedersen FS, Kjems J, Bramsen JB. 2012. Functional selection of shRNA loops from randomized retroviral libraries. PLoS One 7: e43095.

Kaadt E, Alsing S, Cecchi CR, Damgaard CK, Corydon TJ, Aagaard L. 2019. Efficient Knockdown and Lack of Passenger Strand Activity by Dicer-Independent shRNAs Expressed from Pol II-Driven MicroRNA Scaffolds. Mol Ther Nucleic Acids 14: 318–328.

Kampmann M, Horlbeck MA, Chen Y, Tsai JC, Bassik MC, Gilbert LA, Villalta JE, Kwon SC, Chang H, Kim VN et al. 2015. Next-generation libraries for robust RNA interference-based genome-wide screens. Proc Natl Acad Sci U S A 112: E3384–3391.

Knott SRV, Maceli A, Erard N, Chang K, Marran K, Zhou X, Gordon A, Demerdash OE, Wagenblast E, Kim S et al. 2014. A computational algorithm to predict shRNA potency. Mol Cell 56: 796–807.

Kwon SC, Baek SC, Choi YG, Yang J, Lee YS, Woo JS, Kim VN. 2019. Molecular Basis for the Single-Nucleotide Precision of Primary microRNA Processing. Mol Cell 73: 505–518 e505.

Leuschner PJ, Ameres SL, Kueng S, Martinez J. 2006. Cleavage of the siRNA passenger strand during RISC assembly in human cells. EMBO Rep 7: 314–320.

Liu X, Jin DY, McManus MT, Mourelatos Z. 2012. Precursor microRNA-programmed silencing complex assembly pathways in mammals. Mol Cell 46: 507–517.

Liu X, Zheng Q, Vrettos N, Maragkakis M, Alexiou P, Gregory BD, Mourelatos Z. 2014. A MicroRNA precursor surveillance system in quality control of MicroRNA synthesis. Mol Cell 55: 868–879.

Liu YP, Karg M, Harwig A, Herrera-Carrillo E, Jongejan A, van Kampen A, Berkhout B. 2015. Mechanistic insights on the Dicer-independent AGO2-mediated processing of AgoshRNAs. RNA Biol 12: 92–100.

Ma H, Zhang J, Wu H. 2014. Designing Ago2-specific siRNA/shRNA to Avoid Competition with Endogenous miRNAs. Mol Ther Nucleic Acids 3: e176.

Martinez J, Patkaniowska A, Urlaub H, Luhrmann R, Tuschl T. 2002. Single-stranded antisense siRNAs guide target RNA cleavage in RNAi. Cell 110: 563–574.

Matranga C, Tomari Y, Shin C, Bartel DP, Zamore PD. 2005. Passenger-strand cleavage facilitates assembly of siRNA into Ago2-containing RNAi enzyme complexes. Cell 123: 607–620.

Matveeva OV, Nazipova NN, Ogurtsov AY, Shabalina SA. 2012. Optimized models for design of efficient miR30-based shRNAs. Front Genet 3: 163.

McBride JL, Boudreau RL, Harper SQ, Staber PD, Monteys AM, Martins I, Gilmore BL, Burstein H, Peluso RW, Polisky B et al. 2008. Artificial miRNAs mitigate shRNA-mediated toxicity in the brain: implications for the therapeutic development of RNAi. Proc Natl Acad Sci U S A 105: 5868–5873.

Newman MA, Mani V, Hammond SM. 2011. Deep sequencing of microRNA precursors reveals extensive 3’ end modification. RNA 17: 1795–1803.

Okamura K, Liu N, Lai EC. 2009. Distinct mechanisms for microRNA strand selection by Drosophila Argonautes. Mol Cell 36: 431–444.

Paddison PJ, Caudy AA, Bernstein E, Hannon GJ, Conklin DS. 2002. Short hairpin RNAs (shRNAs) induce sequence-specific silencing in mammalian cells. Genes Dev 16: 948–958.

Pelossof R, Fairchild L, Huang CH, Widmer C, Sreedharan VT, Sinha N, Lai DY, Guan Y, Premsrirut PK, Tschaharganeh DF et al. 2017. Prediction of potent shRNAs with a sequential classification algorithm. Nat Biotechnol 35: 350–353.

Premsrirut PK, Dow LE, Kim SY, Camiolo M, Malone CD, Miething C, Scuoppo C, Zuber J, Dickins RA, Kogan SC et al. 2011. A rapid and scalable system for studying gene function in mice using conditional RNA interference. Cell 145: 145–158.

Schopman NC, Liu YP, Konstantinova P, ter Brake O, Berkhout B. 2010. Optimization of shRNA inhibitors by variation of the terminal loop sequence. Antiviral Res 86: 204–211.

Shimizu S, Kamata M, Kittipongdaja P, Chen KN, Kim S, Pang S, Boyer J, Qin FX, An DS, Chen IS. 2009. Characterization of a potent non-cytotoxic shRNA directed to the HIV-1 co-receptor CCR5. Genet Vaccines Ther 7: 8.

Silva JM, Li MZ, Chang K, Ge W, Golding MC, Rickles RJ, Siolas D, Hu G, Paddison PJ, Schlabach MR et al. 2005. Second-generation shRNA libraries covering the mouse and human genomes. Nat Genet 37: 1281–1288.

Starega-Roslan J, Krol J, Koscianska E, Kozlowski P, Szlachcic WJ, Sobczak K, Krzyzosiak WJ. 2011. Structural basis of microRNA length variety. Nucleic Acids Res 39: 257–268.

Sun CP, Wu TH, Chen CC, Wu PY, Shih YM, Tsuneyama K, Tao MH. 2013. Studies of efficacy and liver toxicity related to adeno-associated virus-mediated RNA interference. Hum Gene Ther 24: 739–750.

Vermeulen A, Behlen L, Reynolds A, Wolfson A, Marshall WS, Karpilow J, Khvorova A. 2005. The contributions of dsRNA structure to Dicer specificity and efficiency. RNA 11: 674–682.

Vlassov AV, Korba B, Farrar K, Mukerjee S, Seyhan AA, Ilves H, Kaspar RL, Leake D, Kazakov SA, Johnston BH. 2007. shRNAs targeting hepatitis C: effects of sequence and structural features, and comparision with siRNA. Oligonucleotides 17: 223–236.

Wang Y, Wang YE, Cotticelli MG, Wilson RB. 2008. A random shRNA-encoding library for phenotypic selection and hit-optimization. PLoS One 3: e3171.

Watanabe C, Cuellar TL, Haley B. 2016. Quantitative evaluation of first, second, and third generation hairpin systems reveals the limit of mammalian vector-based RNAi. RNA Biol 13: 25–33.

Wu H, Ma H, Ye C, Ramirez D, Chen S, Montoya J, Shankar P, Wang XA, Manjunath N. 2011. Improved siRNA/shRNA functionality by mismatched duplex. PLoS One 6: e28580.

Xie J, Tai PWL, Brown A, Gong S, Zhu S, Wang Y, Li C, Colpan C, Su Q, He R et al. 2020. Effective and Accurate Gene Silencing by a Recombinant AAV-Compatible MicroRNA Scaffold. Mol Ther 28: 422–430.

Yang JS, Lai EC. 2010. Dicer-independent, Ago2-mediated microRNA biogenesis in vertebrates. Cell Cycle 9: 4455–4460.

Zeng Y, Wagner EJ, Cullen BR. 2002. Both natural and designed micro RNAs can inhibit the expression of cognate mRNAs when expressed in human cells. Mol Cell 9: 1327–1333.

